# The role of deficient action knowledge in unawareness (anosognosia) of tool-action errors after left-hemisphere stroke

**DOI:** 10.64898/2026.03.31.715610

**Authors:** Simon Thibault, Rand Williamson, Aaron L. Wong, Laurel J. Buxbaum

**Affiliations:** Jefferson Moss Rehabilitation Research Institute, Thomas Jefferson University, Elkins Park, PA 19027, USA; Centre de Recherche en Psychologie et Neurosciences, Aix-Marseille Université, 13003 Marseille, France; Department of Rehabilitation Medicine, Thomas Jefferson University, Philadelphia, PA 19107, USA

**Keywords:** Anosognosia, Error Awareness, Limb Apraxia, Action Knowledge, Tool Actions

## Abstract

Many individuals with limb apraxia after left-hemisphere stroke exhibit a lack of awareness of their tool-related action errors, i.e., unawareness of apraxia (UA; also called anosognosia of apraxia). Little is known about the prevalence of UA, the relationship between UA and apraxia severity, or its underlying mechanisms. Here, we assessed both the causes and consequences of UA. Based on a mechanistic model, we hypothesized that UA may arise because of deficits in representations signaling how tool-related movements should look and feel – a component of action knowledge – and that degradation of this knowledge impedes the detection of mismatches between planned and actual tool-related actions. We further predicted that a consequence of UA is a reduction in error-correction attempts. Fifty-six individuals with chronic LCVA gestured to show how to use tools. Immediately after the gesture production task, participants were asked if they made any errors. All participants also completed an action knowledge task to measure the integrity of tool-related movement goals. Individuals were denoted as exhibiting UA if they performed below a normative cutoff for apraxia yet reported making no errors. Our sample included 21 individuals with apraxia; of these, nearly half (48%) exhibited UA. These two groups made a comparable number of gesture errors and were of equivalent stroke severity, yet individuals with UA had significantly more impaired action knowledge. Additionally, individuals with UA were less likely to attempt to correct their errors compared to individuals who were aware of their apraxia. These data are consistent with the hypothesis that action knowledge (how tool actions look and feel) serves a key role in error detection and awareness of apraxia and may contribute to the difficulties with everyday tasks experienced by many people with apraxia.

**Abbreviated Summary:** Thibault et al. show that individuals who are unaware of their tool-use errors (unawareness of apraxia) are selectively impaired at representing how tool-related movements look and feel (action knowledge). Action knowledge may be a central component for the detection of tool-use errors made by individuals with apraxia in everyday tasks.

**Graphical Abstract:** 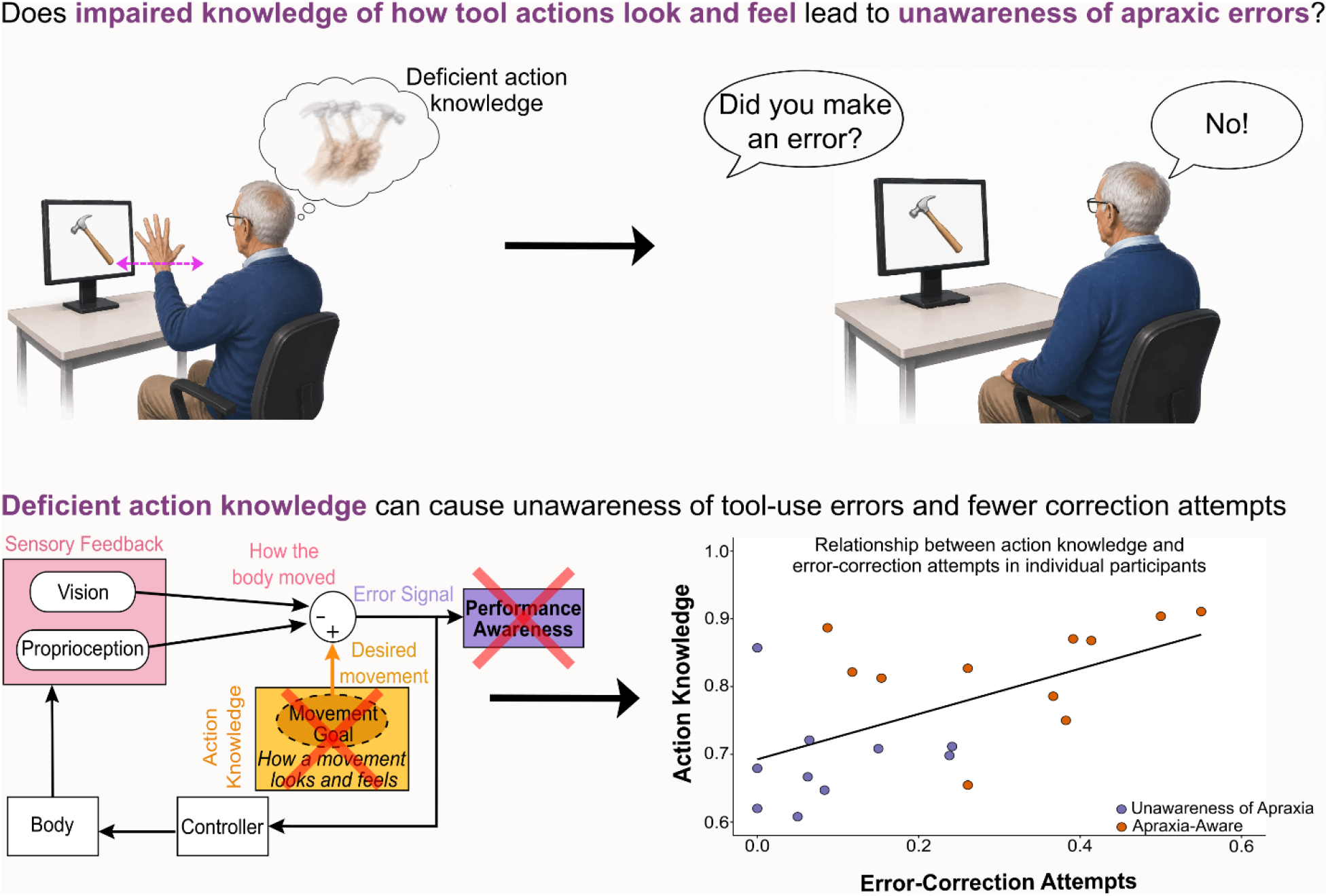

## Introduction

Limb apraxia is a neurological condition that commonly occurs after a left-hemisphere cerebral vascular accident (LCVA – i.e., occlusion or rupture of blood vessels in the brain’s left hemisphere, i.e., a “stroke”) and is characterized by errors in pantomimed tool-use gestures and actual tool use in the absence of elementary sensorimotor deficits^1–7^. Some apraxic individuals exhibit a remarkable phenomenon characterized by unawareness of their praxis deficits (i.e., unawareness of apraxia – UA), a condition also known as anosognosia of apraxia^8–12^. For example, an individual with UA might demonstrate how to use a pair of scissors with a flagrantly incorrect hand posture and movement trajectory but maintain that the gesture was produced correctly when queried about its quality. Deficient awareness of deficit (i.e., anosognosia) has been identified in several other post-stroke syndromes including aphasia, hemiplegia, and spatial neglect, and is linked to poor rehabilitation outcomes and increased caregiver dependence^13^. However, there are very few studies of UA, and little is known about its prevalence or underlying mechanisms. In particular, there is a dearth of knowledge regarding the awareness of deficits in tool-related gestural actions. Previous work on UA has focused on imitation^11,12^, naturalistic actions^8,9^ or composite scores made of different praxis tasks^10^. Given the potential impact of unawareness of for tool-related actions on functional independence, the limited research on this disorder represents a critical gap.

In this study, we tested two hypotheses about the causes and consequences of UA that were based on a novel mechanistic model of error awareness (Fig. 1). Our model has similarities to existing models of performance awareness^14–16^. However, unlike these models, which were developed to address simple movements, our model specifically includes a repository of how skilled praxis actions should look and feel (i.e., action knowledge)^17^. This knowledge serves as a goal for the specification of motor commands sent to the body^2,17–19^ (Fig. 1 - yellow box). In other words, our model is adapted to praxis behaviors, where action knowledge is necessary to inform movement planning. Then, similar to other models, sensory feedback about the movement of the body is sent back to the brain (Fig. 1 - pink box), enabling a comparison between the actual and desired movement. This comparison provides an error signal that facilitates performance awareness; if there is a mismatch between the goal and the produced movement, motor commands are generated to correct the error^20–22^ (Fig. 1 – purple box). Based on this model, our first hypothesis was that reduced action knowledge (and hence an impaired motor goal) gives rise to deficient performance awareness (UA).

**Figure 1:**
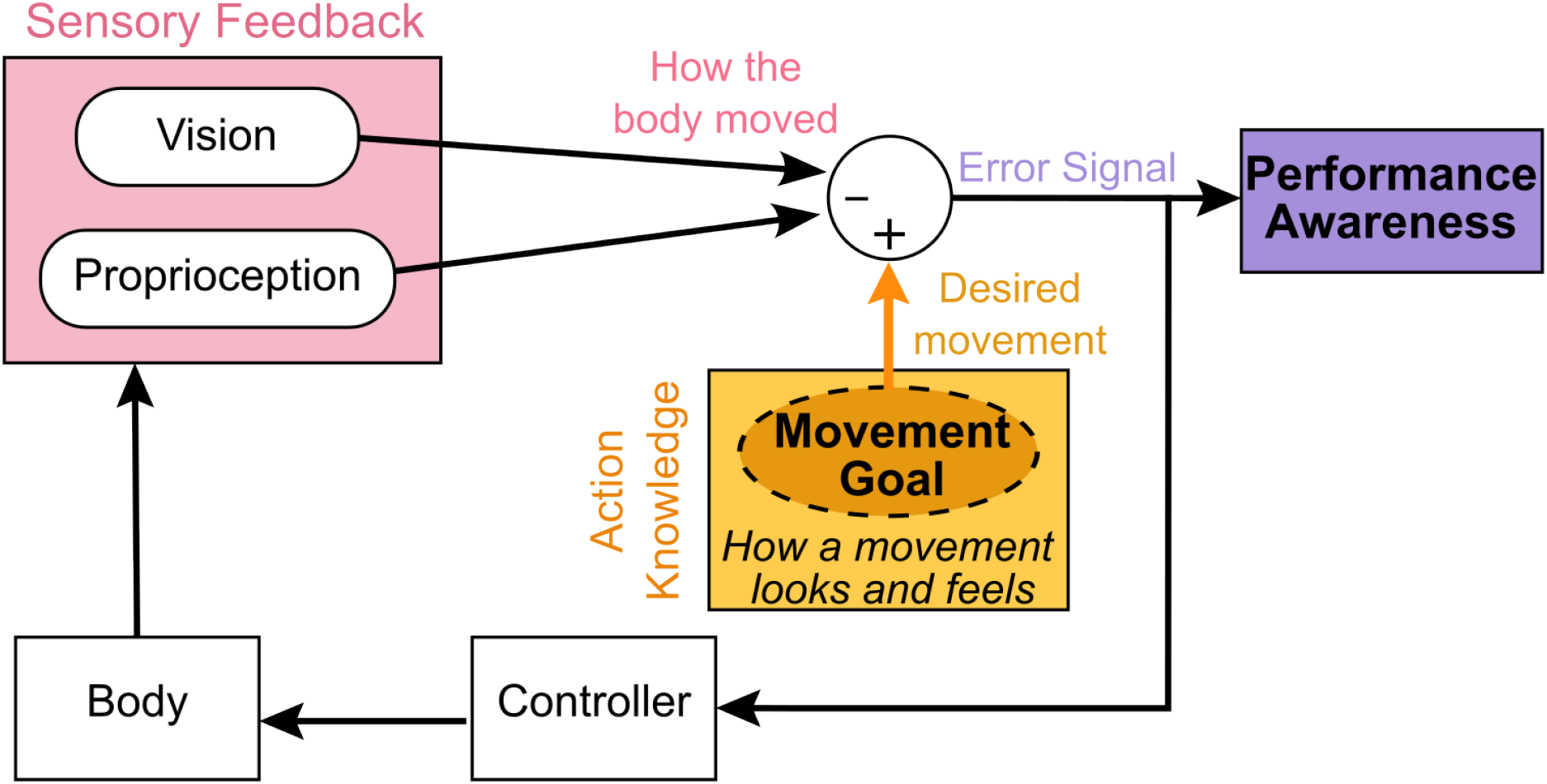
Mechanistic model of performance awareness. Action knowledge serves as a movement goal defining how the movement should look and feel (yellow box). This goal informs motor commands sent to the body. In return, the brain receives visual and proprioceptive feedback that details how the body moved (pink box). This sensory feedback is compared with the desired movement goal, providing an error signal used for performance awareness (purple box). We hypothesize that the movement goal is poorly specified in UA, resulting in a degraded error signal that reduces subsequent error corrections (i.e., via the controller) and increases unawareness of errors.

Our second hypothesis addresses the consequences of UA on the ability to correct one’s errors. Error correction attempts are commonly viewed as a measure of the integrity of performance monitoring and awareness in both language and motor domains^23,24^. In a prior study, we examined error corrections in a tool-use gesture task in individuals with LCVA^17^. We found a positive linear relation between error correction attempts and action knowledge as assessed by a task in which people had to recognize the correct use gesture associated with a given tool^4,17,25^. These data were consistent with the hypothesis that error correction attempts depend upon an intact movement goal. However, UA was not assessed in that study, leaving unclear whether the observed relationship between reduced action knowledge and reduced error correction attempts has any relevance for the clinical UA syndrome. Accordingly, our hypothesis was that we would observe a relationship between action knowledge, error correction attempts, and UA that would not be attributable to the overall severity of neurologic impairment or apraxia.

## Materials and Methods

### Participants

Sixty-six chronic left hemisphere stroke (LCVAs) survivors were recruited from the Research Registry of the Jefferson Moss Rehabilitation Research Institute^26^. All participants experienced a stroke more than 6 months prior to participation, and did not have a history of psychiatric diagnosis, other neurologic disorders, traumatic brain injury, or drug/alcohol abuse. All participants were right-handed prior to stroke onset. To ensure that participants did not have major language comprehension impairments, they were required to achieve a score of 4 or higher (out of 10) on the auditory verbal comprehension subtest of the Western Aphasia Battery^27^ (WAB). Additionally, a measure of stroke severity was acquired for each participant via the National Institutes of Health Stroke Scale^28^ (NIHSS). Hemianopia was assessed with the NIHSS and was relatively rare in the sample (8 individuals). Testing conditions were adjusted for individuals with hemianopic deficits to present stimuli in their intact visual field. Ten of the 66 participants were excluded because they did not have a lesion limited to the left hemisphere (n = 4), were left-handed (n = 3), experienced significant memory or comprehension issues preventing task completion (n = 2) or did not complete all the relevant tasks (n = 1). A sample of 16 right-handed neurotypical adults who did not differ significantly in age, education and sex was also included (Table 1). The neurotypical participants completed a telephone version of the Montreal Cognitive Assessment^29,30^ (MoCA) to confirm that their cognitive functioning was within normal limits (score equal to or above 18). Informed consent was obtained for all participants in accordance with the policies of the Institutional Review Board of Thomas Jefferson University. Participants were compensated for their participation.

**Table 1:** Demographics for the tested sample. To ensure that the groups did not differ significantly in age and education, two-sample t tests were conducted. A chi-squared test was conducted to test whether the proportion of females and males was statistically different across groups. Demographics are reported in the form mean ± standard deviation.

|  | LCVA (N= 56) | Neurotypical Controls (N=16) | Statistics |
| --- | --- | --- | --- |
| **Age (years)** | 63.77 ± 10.56 | 64.69 ± 9.95 | $t_{(70)} = 0.31$; $p = 0.76$ |
| **Education (years)** | 14.50 ± 2.69 | 15.88 ± 2.31 | $t_{(70)} = 1.86$; $p = 0.07$ |
| **Sex** | 24 females | 8 females | $\chi^2_{(1)} = 0.26$; $p = 0.61$ |
|  | 32 males | 8 males |  |
| **Stroke onset (months)** | 115.08 ± 79.74 |  |  |
| **WAB comprehension** | 9.24 ± 0.94 |  |  |
| **Phone MoCA** |  | 20.31 ± 1.20 |  |

### Tasks

#### Gesture-to-Sight (GTS) of Objects Task

Thirty-nine colored photographs of common manipulable objects with a distinct use action were included in the GTS task. These objects included carpentry tools (e.g., hammer), household articles (e.g., clothes iron), office supplies (e.g., scissors), and grooming items (e.g., razor). Participants were seated approximately one meter from a 22-inch computer monitor and instructed to gesture the use of a target object with their ipsilesional left hand. Each object photograph was presented once, in random order, over the course of two blocks. After the object was shown for three seconds, a beep signaled participants to start the gesture. Participants were given 10 s to complete their response; after that time, a “stop” message appeared on the screen and the experimenter instructed them to finish their movement and return to the start position to prepare for the next trial. Gestures were recorded by digital camera and scored offline by one of two trained coders who were reliable with each other (> 85% agreement by Cohen’s Kappa^31^). Each trial was first scored for the presence of a content error (i.e., zero for error, one otherwise) if another recognizable gesture was substituted for the target gesture (e.g., brushing teeth gesture when the object was a comb). Items receiving a content score of one were subsequently scored on three spatiotemporal dimensions: hand posture, arm posture, and amplitude/timing. Following detailed praxis scoring guidelines long in use in our laboratory^25^, each dimension was scored separately and a score of zero was assigned when an error occurred (one otherwise). Occasionally, participants produced multiple gestures; on such trials the best attempt was scored. That is, if participants first produced an error but then adequately corrected the error, they received a score of one for each spatiotemporal dimension. Further coding guidelines were developed to specifically identify any trials with error correction attempts. These trials included a small percentage (8.73%) of trials with successful error correction attempts (i.e., trials on which participants ultimately received the maximum 3-point score). Trials in which the participant produced a content error (0.32% of trials) or stated that they did not recognize the object (1.10% of trials) were removed from the analysis. GTS accuracy was computed as binary score (0 or 1) for each spatiotemporal dimension and each trial. These binary scores were analyzed with statistical models described below. For visualization purpose only, we averaged GTS accuracy for each participant as the average of the three spatiotemporal dimensions (hand posture, arm posture, and amplitude/timing) across all trials.

#### Apraxia error awareness question

We modified an existing questionnaire of unawareness for aphasia^32^ to assess whether participants were aware of their errors. Immediately after the gesture task, they were asked: “During the gesture task, did you ever feel that you knew the purpose or reason for using the object, but found it difficult to show how to use it?”. This wording focuses on any gesture production difficulties participants may have experienced despite good object recognition. This question was asked only a single time after completion of the 39 GTS trials. Participants’ yes or no responses, along with their GTS score, were used to determine whether a participant with apraxia experienced UA (see *Analyses* section for further details). We opted to use this brief assessment because there is no existing measure of unawareness of gestural apraxia. We considered the Visual Analogue Test assessing Anosognosia for Naturalistic Action Tasks (VATA-NAT^8^), but it focuses on naturalistic multi-step actions (e.g., preparing a fried egg), where errors could reflect the influence of capacities other than praxis (e.g. attention control^33^). Moreover, the VATA-NAT is a prospective test measuring the disparity between ratings of a participant and secondary reporter of how well different naturalistic actions would be performed, rather than evaluating an actual attempt to perform the item during the test.

#### Gesture Recognition (GR) Task

The GR task is a verbal action-gesture matching task long in use in our lab^25,34^. Each trial began with a target verb phrase (e.g., “tightening a bolt”) presented on a computer screen along with the simultaneous playing of an audio recording of the phrase. Two short (3-5 second) videos were then played. In one video, an actor was shown gesturing the target action. In the other video, which served as a foil, a spatiotemporal error (29 trials) or semantic error (27 trials) was depicted. Spatiotemporal errors were comprised of target actions with an error in one of the same three dimensions (hand posture, arm posture, or amplitude/timing) used to score performance in the GTS task. Semantic error videos depicted a substitution of an action that was semantically related to the target action. For example, for the target verb phrase “cutting paper”, the foil video depicted a “slicing with a knife” gesture. Participants were instructed to select the video matching the target verb phrase, using the number pad on the keyboard (i.e., “1” for the first video and “2” for the second video). The order of the correct and foil videos was randomized. Stimuli were presented via a program developed on PsychoPy (Open Science Tools Ltd, Nottingham, UK).

Prior to administration of the GR task, we administered a pretest to ensure that individuals could comprehend the verb phrases used in the GR task (e.g., “tightening a bolt”) and knew which tool was associated with that phrase. Participants viewed verb phrases presented singly on a 22-inch computer monitor while an audio recording of the verb phrase was played. Participants then viewed three pictures of numbered tools: the correct tool (e.g., the tool associated with tightening a bolt, i.e., a wrench) and two semantic foils (e.g., a screwdriver and a hammer). Participants were instructed to select, using the number pad on the keyboard, the tool that best matched the verb phrase. There were 34 different test items presented with Eprime 2 (Psychology Software Tools, Pittsburgh, USA). Any verb phrases that a participant failed in the pretest were removed from scoring in the GR task (5.66 % of trials). A subset of the neurotypicals completed the GR task (n = 9). These individuals had an average performance of 0.86 ± 0.06 (performance range: 0.78 – 0.94).

#### NIH Stroke Scale

As a control for overall stroke severity, we administered the NIH Stroke Scale^28^, consisting of 11 items assessing language, consciousness, inattention, vision, motor function, and sensation (Max. score of 42; higher scores indicate greater impairment).

#### Total Lesion Volume

Because NIH Stroke Scale is better suited to assess acute severity rather than chronic severity, we also measured the total lesion volume as complementary index of stroke severity. Total lesion volume was obtained from a subset of participants who were eligible and willing to receive a structural T1 MRI scan (n = 42). Total lesion volume was computed from the lesion maps (voxel size: 1 mm^3^). Lesion maps were either hand-drawn or drawn with an automated algorithm (LINDA^35^). In both cases, the lesion map was checked by an experienced neurologist (H. Branch Coslett) who was blinded to the goals of the study. The lesion maps were registered into the Montreal Neurological Institute (MNI) space.

#### Corsi Block Tapping Task (working memory)

As as control for working memory deficits, we administered the Corsi task^36^ in PEBL^37^ to assess visuo-spatial memory span. Participants were presented with rectangles on a computer screen. On each trial, rectangles would momentarily light up one by one in a particular sequence. At the end of the sequence, participants used the computer mouse to click the rectangles in the same order that they were presented. The sequence length started at 2 rectangles and increased by 1 each time the sequence was correctly reproduced; participants had 2 attempts to reproduce a sequence of a given length. The task ended when the participant failed both attempts; the total step length successfully completed was reported as the memory span.

### Data Analyses

#### Identifying individuals with apraxia and UA

LCVA participants were subdivided into apraxic and non-apraxic groups based on a cutoff score calculated from the average GTS scores across all trials of the neurotypical participants. LCVA individuals scoring less than 0.85 (i.e., more than two standard deviations below the mean of the neurotypicals) were classified as apraxic. People with apraxia were further subdivided according to their responses to the apraxia awareness question. If participants with apraxia denied that they had issues showing how to use tools in the GTS task, they were classified as having UA. The LCVA sample was thus subdivided into three groups: non-apraxic individuals (Non-Apraxic), individuals experiencing apraxia with awareness of their deficit (Apraxic-Aware), and individuals experiencing unawareness of apraxia (UA).

#### Statistical analyses

To test the study predictions, we analyzed GTS and GR accuracy, as well as the proportion of trials with error correction attempts in the GTS task. We also considered whether UA was related to apraxia severity and/or overall stroke severity using GTS accuracy and the NIHSS score, respectively. To estimate the effect of others potential confounds, we also analysed total lesion volume, time since stroke onset, working memory span, and aphasia severity (WAB comprehension subset). GTS accuracy, GR accuracy and error-correction attempts were analyzed as trial-level data, where each participant had many observations. GTS accuracy, GR accuracy and error correction attempts were binomial scores (0 or 1 at the trial-level); thus, we modeled them using Generalized Linear Mixed Models (GLMM) with binomial response distributions. We averaged the trial-level data (GTS Accuracy, GR accuracy and error-correction attempts) for each participant for visualization purposes (see Figures 2 and 3). In contrast, NIHSS scores consisted of only one observation per participant, so we analyzed this score with a one-way ANOVA. Likewise, the four potential confounds noted earlier were analyzed with one-way ANOVAs.

**Figure 2:**
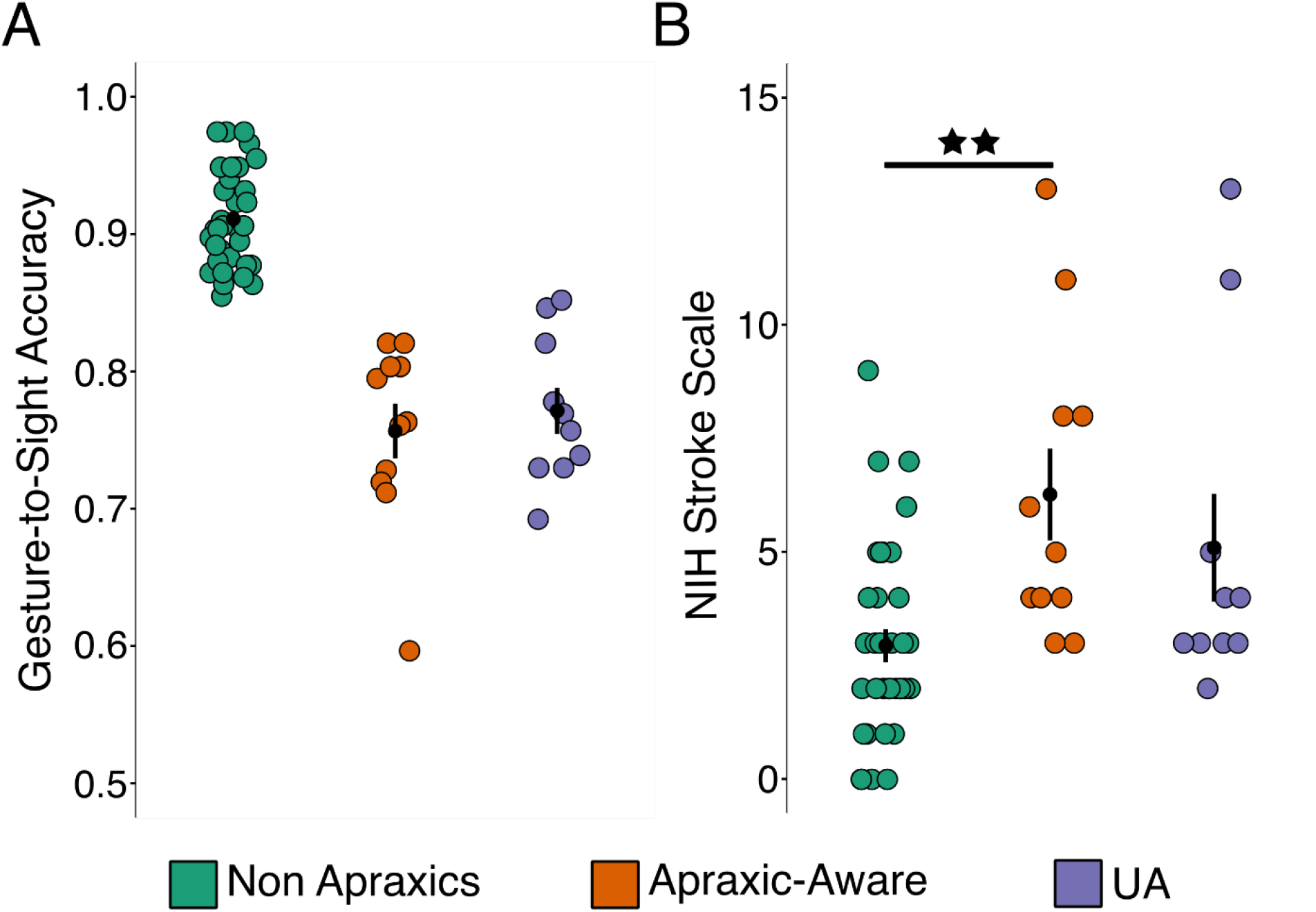
Individuals with UA showed equivalent performance to other groups in apraxia and stroke severity. A) Plot shows the proportion of correct trials on the Gesture-to-Sight (GTS) task assessing apraxia severity. The twenty-one apraxic participants were tested with a GLMM on GTS accuracy. No difference in GTS accuracy was observed between Apraxic-Aware [0.76 ± 0.02] and UA [0.77 ± 0.02] groups (χ^2^_(1)_ = 0.32, p = 0.57). B) Plot shows stroke severity (NIH Stroke Scale) between the three groups [Non-apraxic = 2.94 ± 0.36; Apraxic-Aware = 6.27 ± 1.10; UA = 5.10 ± 1.19; F_(2, 53)_ = 7.21, p = 0.002]. Fifty-six participants were tested with a one-way ANOVA. Each colored dot represents a unique participant. The group average is depicted by black dots, and error bars represent the group standard error of the mean (SEM). Significance level is indicated with a star: ★★ p < 0.01.

**Figure 3:**
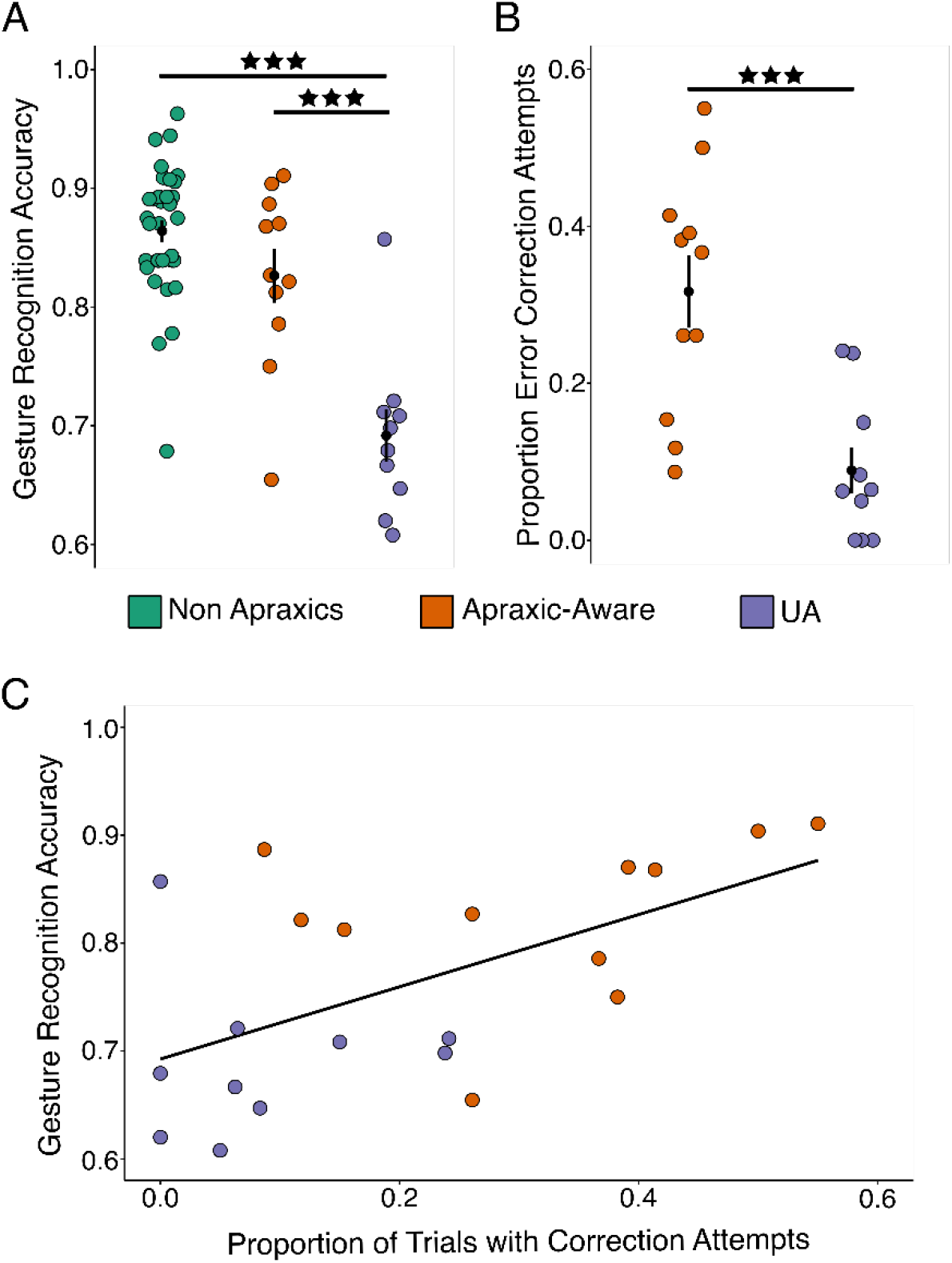
Individuals with UA had more deficient action knowledge and fewer error correction attempts than individuals in the Apraxic-Aware group. A) Individuals with UA showed a selective impairment in gesture recognition (GR) when compared to the other two groups [Non-apraxics = 0.86 ± 0.01; Apraxic-Aware = 0.83 ± 0.02; UA = 0.69 ± 0.02; χ^2^_(2)_ = 35.02, p < 0.001]. Fifty-six participants were tested with a GLMM on GR accuracy. B) Proportionally fewer error trials were associated with correction attempts in UA than Apraxic-Aware individuals. [Apraxic-Aware = 0.32 ± 0.05; UA = 0.09 ± 0.03; χ^2^_(1)_ = 13.71, p < 0.001]. The twenty-one apraxic participants were tested with a GLMM on error-correction attempts. The group average is depicted by black dots, and error bars represent the group standard error of the mean (SEM). Significance levels are indicated with stars: ★★★ p < 0.001. C) The correlation between GR accuracy and the proportion of trials with error corrections was significantly positive (r = 0.58; p = 0.003). The twenty-one apraxic participants were tested with a Pearson correlation. Each colored dot represents a unique participant.

To assess whether the Apraxic-Aware and UA groups differed in GTS accuracy, we included Group as a fixed effect. By definition, these groups had lower GTS accuracy than the Non-Apraxic group, so the Non-Apraxic group was not included in this analysis. Participant was included as a random effect. NIHSS data were analyzed with group as a fixed effect (Non-Apraxic vs. Apraxic-Aware vs. UA). For GR accuracy, we included Group (Non-Apraxic vs. Apraxic-Aware vs. UA) as a fixed effect and Participant as a random effect. For the proportion of trials with error correction attempts, we use a similar model but only included the Apraxic-Aware and UA groups (as these groups produced sufficient errors for analysis). Where relevant, Tukey post-hoc tests were conducted to compare pairwise differences between groups. Finally, because we had an *a priori* directional hypothesis that action knowledge would be positively associated with error-correction attempts, we ran a one-sided Pearson correlation between total GR accuracy and the proportion of GTS trials with error correction attempts. All analyses were conducted in R 4.3.3 (R Core Team). The package *afex 1.3-1* was used for statistical analyses and the package *emmeans 1.10.1* for posthoc tests.

## Results

### UA is common in individuals with apraxia

Of the 56 LCVA individuals tested, 21 had limb apraxia and 35 were Non-Apraxic. Among the individuals with apraxia, there were 10 individuals with UA and 11 Apraxic-Aware individuals. Thus, in our sample 47.62% of individuals with apraxia experienced UA. The proportion of individuals with UA is broadly consistent with a previous study in which 50% of individuals with limb and/or bucco-facial apraxia showed reduced awareness of their deficit^10^.

### Apraxia severity and overall stroke severity are not associated with UA

We first tested whether UA could simply be explained by the severity of apraxia or overall cognitive and sensory-motor deficits related to stroke. There was no difference in the performance of the Apraxic-Aware versus UA groups on the GTS task (χ^2^_(1)_ = 0.32, p = 0.57; Fig. 2A). For the NIHSS, although we found an effect of group (F_(2, 53)_ = 7.21, p = 0.002; Fig. 2B), post-hoc tests revealed that this was not attributable to differences between UA individuals and either the Apraxic-Aware (p = 0.81) or Non-Apraxic (p = 0.08) groups. Rather, Apraxic-Aware individuals had significantly greater overall stroke impairment relative to Non-Apraxic individuals (p = 0.003). To confirm this finding, we analyzed total brain lesion volume in the subset of individuals who received a brain scan (n = 42); no significant group effect was observed (F_(2, 39)_ = 2.49; p = 0.10; Non-Apraxic = 59.58 ± 11.07 cm^3^; Apraxic-Aware = 113.22 ± 17.99 cm^3^; UA = 86.33 ± 26.10 cm^3^). Together, these findings suggest that UA is explained neither by apraxia severity nor overall stroke severity. We also assessed whether the groups differed in time since stroke onset, working memory, or aphasia severity (WAB comprehension). There was no significant difference between groups for time since stroke onset (F_(2,53)_ = 1.98; p = 0.15; Non-Apraxic = 99.00 ± 13.01 months; Apraxic-Aware = 138.93 ± 21.23 months; UA = 145.15 ± 28.77 months) or working memory (F_(2,53)_ = 1.67; p = 0.20; Non-Apraxic = 4.49 ± 0.13; Apraxic-Aware = 3.95 ± 0.30; UA = 4.50 ± 0.32). There was a group difference for aphasia severity (F_(2,53)_ = 6.02; p = 0.004; Non-Apraxic = 9.53 ± 0.11; Apraxic-Aware = 8.97 ± 0.32; UA = 8.52 ± 0.39), but importantly the difference between UA and Apraxic-Aware individuals was not significant (p = 0.45). Rather, aphasia severity was greater for individuals with UA compared to Non-Apraxic individuals (p = 0.005). The Apraxic-Aware and Non-Apraxic individuals did not differ in aphasia severity (p = 0.16).

### Individuals with UA have a selective impairment in action knowledge and are less likely to correct errors

To test the hypothesis that UA is caused by deficient action knowledge, we analyzed Gesture Recognition (GR) accuracy across all three groups. We found a group effect (χ^2^_(2)_ = 35.02, p < 0.001); post-hoc tests revealed an impairment in action knowledge for people with UA when compared to both the Apraxic-Aware (p < 0.001) and Non-Apraxic groups (p < 0.001; Fig. 3A). No statistical difference was observed between the Non-Apraxic and Apraxic-Aware groups (p = 0.15). Viewed together with the results reported above, this indicates that individuals with UA have a specific deficit in action knowledge.

Our second prediction was that UA would impact individuals’ abilities to detect and correct their own errors. We found that the proportion of error trials that contained an error correction attempt was significantly smaller for individuals with UA compared to Apraxic-Aware individuals (χ^2^_(1)_ = 13.71, p < 0.001; Fig. 3B). Moreover, we found that action knowledge (GR accuracy) was positively correlated with the proportion of error trials with error correction attempts (r = 0.58; p = 0.003; Fig. 3C). Thus, individuals with poor action knowledge are less likely to correct their errors.

## Discussion

Limb apraxia is a common disorder, occurring in 25%-50% of individuals following a stroke in the left hemisphere^38,39^ and is associated with poor post-stroke recovery^40^. A significant proportion of these individuals (nearly 50% in our study) are not aware of their impairments, suggesting that UA may be a major contributing factor to disability. Why UA arises and how it impacts tool-use ability has not been well-understood. This study tested two predictions derived from a mechanistic model of performance awareness (Fig. 1) in which action knowledge serves as a movement goal that specifies how an action should look and feel^2,17–19^. According to this model, imprecise and/or slowly-activated action knowledge in individuals with UA disrupts the ability to compare movement goals against executed movements, with resulting imprecision or “noise” in an error “mismatch” signal that enables error detection and correction. Thus, we hypothesized that UA should be specifically associated with both deficient action knowledge and reduced error correction attempts. The findings of the current study are consistent with these predictions and additionally suggested a reliable relationship between the magnitude of impairment in action knowledge and the magnitude of reduction in error correction attempts.

It is noteworthy that we found no relationship between UA and apraxia severity, consistent with the results of a prior study^10^. This suggests that UA does not simply reflect an overall degradation of the praxis system. Instead, our data suggest that, at least in some individuals, UA is related to a deficit in action knowledge (i.e., stored movement goals). Note that not all apraxic impairments arise from action knowledge deficits; they may also arise from a downstream disruption in transforming movement goals into a motor command to be executed^2,3,41–46^. The locus of impairment could give rise to different patterns of behavior: while impairments at the level of action knowledge would impact the ability to recognize one’s own errors as well as others’ errors, downstream disruptions would likely only disrupt the former. Gesture recognition tasks can be used to probe action knowledge impairments, whereas tests of gesture imitation could be used to probe downstream impairments. This is because imitation does not necessitate accessing stored action knowledge; instead, visual input (i.e., the gesture to be imitated) can be directly transformed into a representation that guides the desired movement^47^. Indeed, a prior study using action imitation to probe UA at this more downstream point along the praxis pathway found an association between apraxia severity and the ability to detect self-produced (but not other-produced) errors^12^. Taken together with the present results, these data suggest that damage at different loci may contribute to unawareness of tool-use production errors (where a stored movement goal is required) versus imitation errors (where one’s own actions must be matched to an action recently performed by another person, and where a failure to detect mismatches could result in unawareness).

Consistent with this possibility, our model suggests several possible loci for unawareness of praxis performance errors. First, impairments in sensory feedback processing (Fig. 1 – pink box), may result in deficient awareness that generated movements deviate from movement goals. We recently demonstrated that individuals with apraxia exhibit a specific reduction in the ability to use proprioceptive information to guide movements^18^. Accordingly, it is possible that UA in some individuals may be attributable to deficits in using proprioceptive feedback to detect their own errors. Alternatively, performance unawareness may arise from an impaired ability to compare intact movement goals to generated movements, as has been hypothesized for anosognosia of hemiplegia^14,15,48^. Both the proprioceptive feedback and “comparator” accounts could explain the disparity between impaired detection of one’s own errors and intact detection of other errors that was observed by Scandola and colleagues (2021) in an imitation task. In summary, error monitoring failures may plausibly arise for at least three reasons: failure to retrieve a movement goal, deficits in proprioceptive feedback integration, or deficient comparison of intact goals with feedback. Furthermore, the locus of failure may depend in part on the input and output requirements of the task, as well as other cognitive processes such as attention and memory. While our work was primarily dedicated to understanding the role of deficits in action knowledge, it will be of interest to investigate these different components in future research to uncover how they may contribute altogether to UA.

Error correction attempts by apraxic-aware individuals suggest that they were able to detect the discrepancy between the expected and actual movement after the movement was produced, although they were unable to prevent the error from occurring in the first place^17^. In other domains (e.g., speech, typing), error correction attempts are commonly viewed as a key index of the integrity of performance monitoring and a window into the processes that are used for error detection^49^. In picture naming, correction attempts consisting of successive approximations of the target word (“conduite d’approche”) are a hallmark of conduction aphasia, a language disorder characterized as a failure to transmit intact knowledge of phonemes/syllables/words to an intact production planning system in a timely way to prevent errors. In this disorder, errors are detected by the auditory comprehension system as a mismatch between the way the phoneme/syllable/word is supposed to sound and auditory feedback about what was actually produced. These detected errors in turn prompt another attempt at producing the correct utterance: approximations to the target word are thus viewed as “clean-up” attempts by the intact comprehension system^50–52^. Indeed, in several language models^53^, intact knowledge of the movement goal (how an utterance should sound) is a prerequisite for error correction^54,55^. We propose that apraxic-aware individuals leverage their spared action knowledge (i.e., knowledge that is equivalent to that of non-apraxic individuals) to employ a similar correction strategy based on the desired look and feel of tool actions. Yet these individuals still experience apraxia, which may be explained by their inability to select the appropriate action among competitors^56–58^, or their incapacity to integrate multisensory feedback during action planning^18^. As noted earlier, many classic and contemporary accounts of apraxia posit two major subtypes: one characterized by deficits in action knowledge (ideational apraxia), and the other by a disconnection between intact knowledge and motor planning (ideomotor apraxia)^2,3,41–46^. The former subtype, typically referred to as ideational or conceptual apraxia, is described as an inability to conceptualize a proper action plan^59,60^. The term ideational apraxia dates to Liepmann^43^, who speculated that it reflects a loss of action concepts (i.e., the idea of an action). Similarly, Morlaas^61^, DeRenzi^59^, and Ochipa and colleagues^62^ characterized ideational apraxia as a specific loss of action knowledge. Our view is that ideational apraxia is most clearly identified by poor performance on gesture recognition or other concept-focused tasks that do not have a gestural output requirement, such as those tapping manipulation knowledge^63^. In contrast, ideomotor apraxia has been linked to underlying deficits in processes further “downstream”, such as the integration of sensory information to form a motor plan^18^ or selecting among action competitors^56–58^. In its purest form, ideomotor apraxia is associated with spatiotemporal errors in production in the context of intact gesture recognition^64^. Nevertheless, differential diagnosis is often difficult: gesture errors are produced in both subtypes for different reasons, and action recognition and production deficits often co-occur to varying degrees^2,3,65,66^. Our work tested the novel hypothesis that UA is related to loss of action knowledge (putatively, the core deficit in ideational apraxia). This relationship is not trivial, as individuals could theoretically lose knowledge of actions and be well-aware that their performance is degraded. For instance, in neurotypical individuals, being unable to recall a fact that one previously knew is typically associated with awareness that one has forgotten that fact. The relationship we have observed here suggests instead that loss of action knowledge is closely tied to loss of praxis error awareness. Given the strong relationship we observed between action knowledge recognition and the ability to detect one’s own errors, it may be clinically useful when attempting differential diagnoses to consider poor error detection and UA as hallmarks of the ideational apraxia syndrome, whereas apraxia with error awareness may be more consistent with a diagnosis of ideomotor apraxia.

The present study had several limitations. There are existing anosognosia questionnaires for hemiplegia^67^, spatial neglect^68,69^, and everyday complex actions^8,9^, but we are not aware of a specific assessment for gestural praxis actions. Our UA assessment, which we developed based on a prior questionnaire for assessing unawareness of aphasia^32^, was very brief and administered only at the end of the GTS task. On the other hand, we found a strong correspondence between this end-of-task measure and a trial-level measure in the subset of participants (n = 42) for whom we had data on both measures. Individuals with UA were less likely to detect their errors compared to apraxic-aware individuals when their awareness was assessed on a trial-level basis (χ^2^_(2)_ = 11.63; p = 0.003; Apraxic-Aware = 37.49 ± 8.09% vs. UA = 6.77 ± 3.52%). Nevertheless, future studies would benefit from including a trial-level assessment for all participants. Moreover, the temporal dynamics of perception of performance may be an interesting issue for future research. For example, it may be possible that some individuals can immediately judge their performance on a given trial as impaired but be unable to integrate this into a long-term belief that they have problems with the task as a whole^70–72^. Another potential limitation of our study is that chronic stroke severity was assessed with the NIH Stroke Scale, which may be relatively insensitive in chronic stroke. Nonetheless, the lack of confounding influence of stroke severity was confirmed with an analysis of total lesion volume, which also did not differ across the three groups. Finally, from both theoretical and clinical perspectives it will be of interest to further examine the relationship of UA to anosognosia in other domains (e.g., hemiparesis, aphasia, spatial neglect), as well as to further explore the hypothesis that UA may arise because of deficits at several different points in processing (e.g., sensory integration or comparison), as noted earlier. Each point in processing is dependent on the praxis ability considered (i.e., imitation vs. gesture) and whether unawareness deficits are consistent across praxis abilities need to be clarified. Finally, the role of attention, memory and executive functions to unawareness deficits will also need to be characterized more precisely in future studies.

Despite these limitations, the results we report may be informative from a clinical perspective. Treatments for limb apraxia often focus on practicing gesture production^73^. The present findings suggest that rehabilitation approaches may benefit from an additional focus on restoring action knowledge in individuals with UA. Indeed, prior research indicates that remediation of action knowledge may also benefit gesture production accuracy^74^. Assessing UA in rehabilitation settings may facilitate the provision of interventions targeted at individuals’ reduced self-awareness and help to improve caregivers’ awareness of this challenge. Taken together, our findings of the relationship of UA to impaired action knowledge and reduced error correction highlight the importance of systematically assessing UA when testing for the presence of limb apraxia.

## Data Availability

Relevant data and scripts are available at https://doi.org/10.5281/zenodo.21980049.

## Acknowledgments

We are grateful to John Yates for his support in recruitment, acquisition, and data preprocessing.

## Funding

This work was supported by NIH Grant #R01NS115862 to Aaron L. Wong.

## Competing Interest

The authors have no competing interest to declare.

## Notes

### Competing Interest Statement

The authors have declared no competing interest.

### Summary of Updates

Minor changes to report some averages based on a reviewer's request

